# Acriflavine delivery via Polyurethane nanocapsules to treat neovascular age-related macular degeneration

**DOI:** 10.64898/2025.12.22.695712

**Authors:** Narendra Pandala, Lorena De Melo Haefeli, Adnan Khan, Hailey Steffan, Jake Miller, Edwin M. Stone, Ian C. Han, Erin B. Lavik, Robert F. Mullins, Budd A. Tucker

## Abstract

Choroidal neovascularization is a complication associated with retinal diseases such as age-related macular degeneration (AMD), a leading cause of vision loss in the developed world. Choroidal neovascular membrane (CNVM) refers to the abnormal growth of blood vessels in the retina which results in exudation and/or hemorrhage, leading to photoreceptor damage and vision loss. Currently first-line treatment for CNVM include intravitreal injections of vascular endothelial growth factor (VEGF)-binding antibodies that prevent the growth of these leaky blood vessels. Unfortunately, anti-VEGF drugs often require frequent injections, and prolonged VEGF inhibition has been associated with retinal atrophy and decreased long term effectiveness in some patients. This study presents the use of Acriflavine, a small molecule HIF1α inhibitor loaded polyurethane nanocapsules to treat CNVM in a rat model. Fourteen days following laser injury and intravitreal drug administration, CMVM size was significantly reduced in acriflavine nanocapsule and free acriflavine treated animals as compared to drug free controls. Moreover, acriflavine nanocapsules reduce CNVM incidence compared to drug free controls by approximately 25%. Among the different delivery routes tested, intravitreal delivery of acriflavine nanocapsules was found to be superior to subretinal and suprachoroidal delivery for reducing CNVM area without causing significant damage to the neural retina. This paper presents the synthesis, characterization and the effectiveness of the polyurethane based acriflavine delivery system in treating choroidal neovascularization.

## Introduction

Age-related macular degeneration (AMD) is a blinding retinal degenerative disorder that affects millions of individuals worldwide. While most AMD patients are diagnosed with the slowly progressive atrophic form of the disease (i.e., dry AMD), neovascular AMD (nAMD), which accounts for approximately 10-15% of patients (*1*), is characterized by aberrant growth of leaky blood vessels and buildup of fluid in the central region of the retina known as the macula (*2*). These abnormal blood vessels in most cases originate from the choroid, extend into the retina through Bruch’s membrane, and form a dense vascular network either beneath the retinal pigment epithelium (RPE) or in the subretinal space. When they arise from the choroidal vasculature, these pathologic vascular lesions are referred to as choroidal neovascular membranes (CNVMs) (*2*). CNVMs are frequently associated with subretinal fluid leakage and retinal fibrosis, resulting in rapid vision loss (*3*). In addition, inflammation (*4*), polarization of macrophages towards proangiogenic phenotypes (*5*), and oxidative stress (*6*) have been reported in nAMD.

Current nAMD treatments are focused on reducing angiogenesis via intravitreal injection of antibodies targeting Vascular Endothelial Growth Factor (VEGF) (*7–12*). VEGF is a key regulator of angiogenesis that has been studied extensively in the context of cancer and more recently ocular disease. For instance, several inhibitory anti-VEGF antibodies are now in clinical use for the treatment of nAMD (*7–12*). While anti-VEGF antibodies are highly effective, they are short lived and require re-administration as often as every four weeks. Repeated intravitreal injections carry the risk of ocular complications such as cataracts and endophthalmitis, and place a significant care burden on elderly patients, many of whom can no longer drive (*13, 14*). Although long-term formulations are desirable, delivery systems have been difficult to develop (*15*). In addition, as VEGF plays an important role in maintaining health of the neural retina (*16*), persistent inhibition may have detrimental effects on photoreceptor cell health, potentially contributing to the development of geographic atrophy (*17*). For these reasons, alternate therapeutic targets with more durable treatment effects are highly desirable.

Hypoxia-inducible factor 1 (HIF1) is an oxygen sensitive transcriptional activator that functions as a master regulator of angiogenesis (*18, 19*). This heterodimeric complex consists of an α and *β* subunit. While the HIF-1*β* subunit is constitutively expressed and abundantly available, HIF-1α is translated but continuously degraded via the ubiquitin pathway under normoxic conditions (*18, 20, 21*). Under hypoxic conditions, HIF-1α degradation is arrested in the cytoplasm and the subunit is in turn translocated to the nucleus where it binds HIF-1*β* and regulates expression of angiogenic factors, glucose transporters, and glycolytic enzymes (*18, 20, 21*). Because loss of HIF-1α in endothelial cells has been shown to inhibit VEGF induced angiogenesis (*22*), we believe it is an ideal target for nAMD treatment.

Acriflavine, a small molecule used as a topical antiseptic, has recently been identified as an effective HIF-1α inhibitor capable of mitigating ocular neovascularization in animal models (*23–27*). However, due to its small size, it is quickly cleared from the eye following intravitreal injection (*23, 26*), limiting its potential for clinical application. Unlike anti-VEGF antibodies, acriflavine is exceptionally stable and has been incorporated into several different drug delivery formulations (*26, 28, 29*). For instance, Hackett and colleagues demonstrated successful loading of acriflavine into PLGA microparticles and release over a 60-day time frame *in vitro* (*26*). They went on to demonstrate long term suppression of CNVM formation via suprachoroidal versus intravitreal injection in a mouse laser injury model (*26*). As promising as these studies are, PLGA degrades quite quickly and in large doses can cause an acidic PLGA response(*30–32*). In addition, up to 90% of the drug is released from PLGA microparticles as a single burst within the first 1-2 weeks following delivery.

To address these issues, we recently reported development of a novel polyurethane nanocapsule formulation capable of both sustained acriflavine release and ultrasound mediated on-demand delivery *in vitro* (*29*). In this study our goal was to determine if ocular delivery of acriflavine loaded polyurethane nanocapsules was safe and capable of mitigating CNVM formation in a rat model of laser injury *in vivo* (**Figure 1A**).

**Figure 1.**
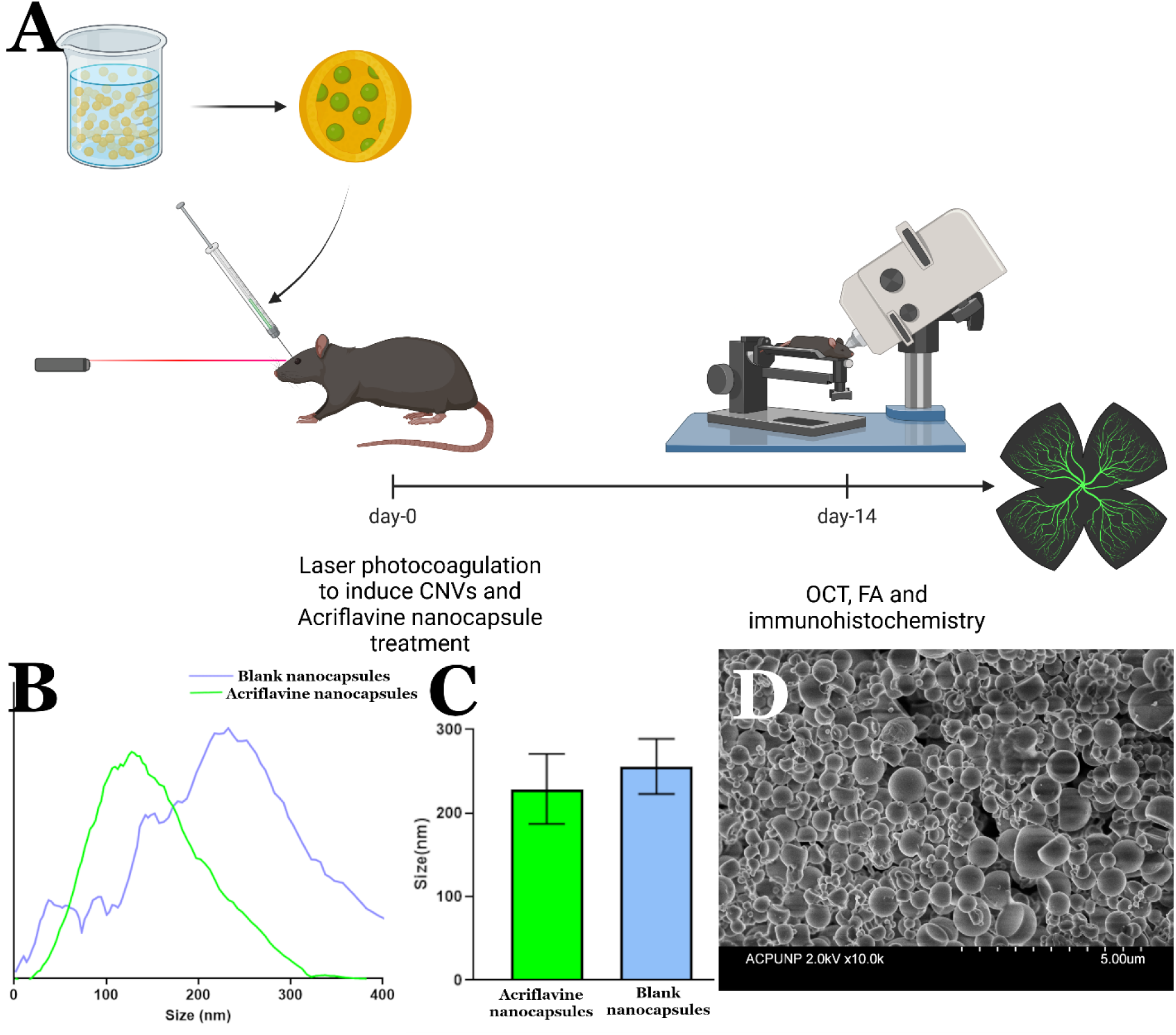
Schematic of synthesis and Acriflavine nanocapsule testing. A) Schematic showing the Polyurethane nanocapsules loaded with Acriflavine administered in a rodent model of laser induced CNV. B) Representative size distribution of the Acriflavine loaded and the blank nanocapsules. C) Average size of the nanocapsules (n=3 preparations) D) Scanning electron microscopy (SEM) images showing the morphology of the Acriflavine loaded polyurethane nanocapsules.

## Results

### Synthesis and characterization of the nanocapsules

Polyurethane nanocapsules were synthesized as previously described using an interfacial condensation polymerization reaction in an oil in water nano emulsion (*29*). Following fabrication, nanocapsules were coated with polyethylene glycol (PEG) and poloxamer to increase hydrophilicity and make it easier to generate injectable single particle suspensions. Polyurethane nanocapsules without drug (blank nanocapsules hereafter) have a size of 256±33 nm while acriflavine loaded polyurethane nanocapsules (acriflavine nanocapsules hereafter) have a size of 229±42 nm (**Figure 1B**). The drug loading was found to 49±3 μg of acriflavine per mg of nanocapsules, with an encapsulation efficiency of ∼12%. Scanning electron microscopy (SEM) demonstrating the size range and distribution of nanocapsules present (**Figure 1C**).

### In vitro testing

To test efficacy and biocompatibility of acriflavine nanocapsules *in vitro*, we first performed a scratch assay using our previously published immortalized human choroidal endothelial cell line ICEC2-TS (*33, 34*) (**Figure 2**). After 24 hours, we observed that both the free drug and acriflavine nanocapsules (at 10 µg/ml and 20 µg/ml) prevented the regrowth of cells into the scratch area compared to the controls, without any signs of cytotoxicity (the cells outside the scratch area remained unchanged) (**Figure 2**).

**Figure 2.**
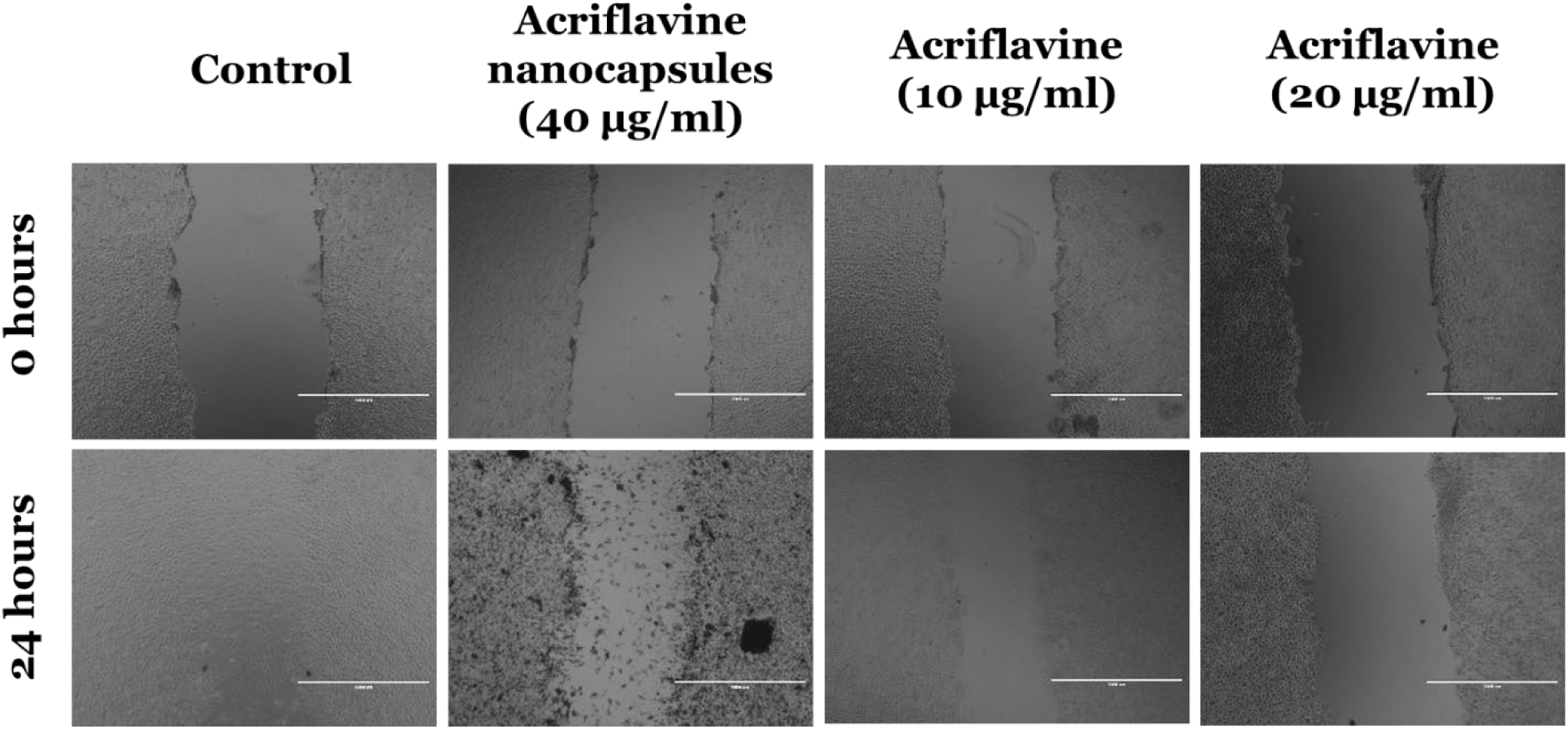
Acriflavine inhibits endothelial cell migration *in vitro*. Wound healing assay using a human choroidal endothelial cell line showing the effect of free acriflavine and acriflavine nanocapsules in preventing wound closure after 24 hours (Scale bar is 1000 µm; This is an immortalized cell line generated from primary human choroidal endothelial cells).

### Intravitreal acriflavine without nanocapsules reduces CNVM size

To evaluate efficacy of Acriflavine *in vivo,* the rat laser photocoagulation model of choroidal neovascularization was employed. As per the methods section, free acriflavine (3 ug/ul; 10 ul per eye) was injected intravitreally immediately following laser injury. Animals were analyzed at 14-days following treatment using fluorescent fundus angiography, optical coherence tomography, and immunohistochemistry of RPE/choroid flat mounts. CNVMs were readily identifiable in all three imaging modalities (**Figure 3A-C**). Fundus angiography and OCT imaging provided clear images with no signs of inflammation or cataract formation in any animal (**Figure 3A, B**). CNVMs were smaller in Acriflavine treated eyes as compared to drug free control eyes (**Figure 3D**).

**Figure 3.**
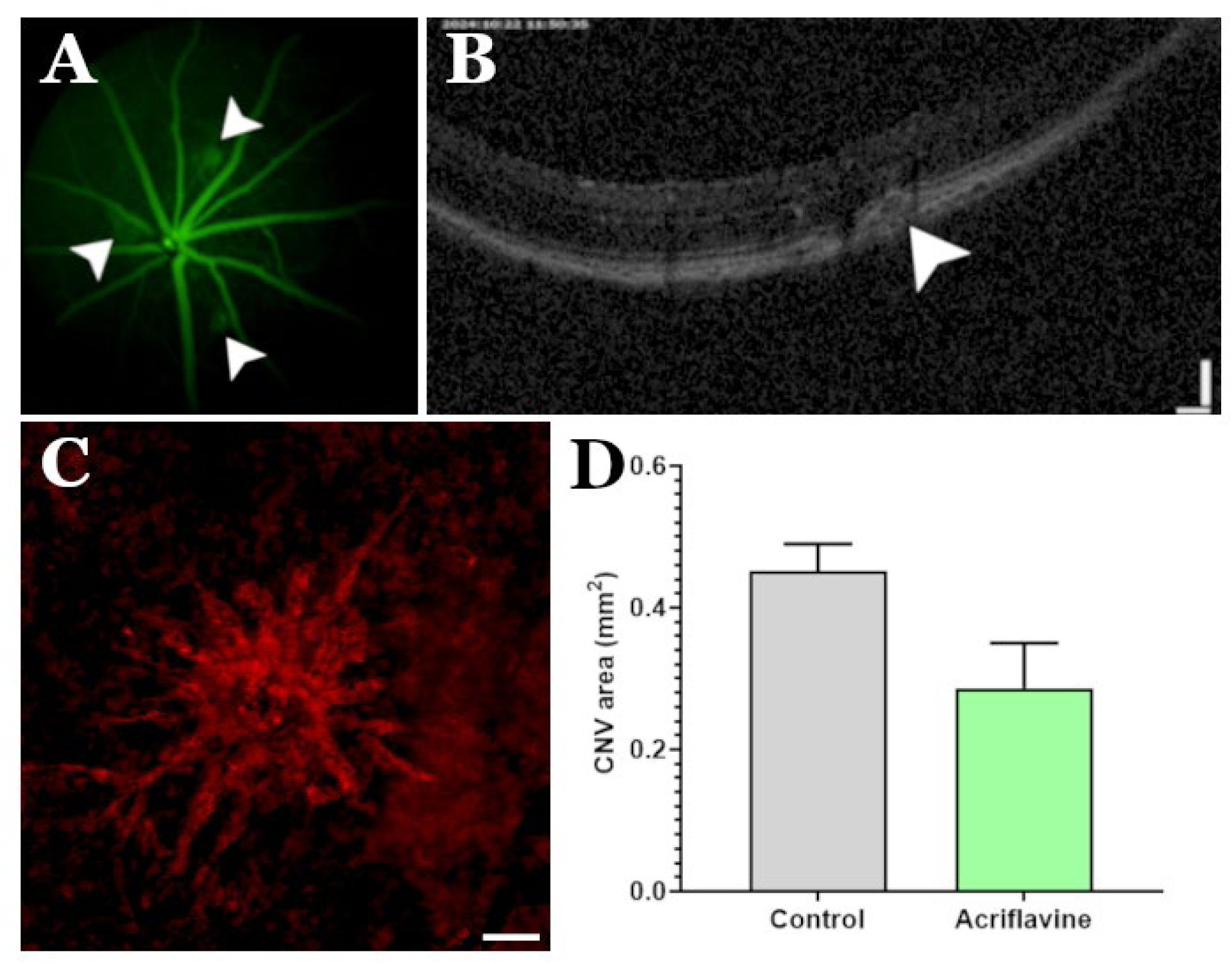
Acriflavine mitigates CNVM formation *in vivo*. A) Fluorescent fundus angiography demonstrating the presence of laser induced CNVMs in the eye of a Brown Norway rat at 14-days post-procedure (white arrows). B) Optical coherence tomography (OCT) image showing the presence of the CNVM (white arrow) at the lesion site and lack of vitreous inflammation. C) GSL-1 ((*Griffonia simplicifolia* lectin I) labeling showing the CNVM in a flat mount preparation of the RPE-choroid complex (Scale bar is 1000 um). D) Analysis of CNVM area demonstrating the effect of Acriflavine *in vivo* (delivered via an intravitreal injection).

### Impact of Acriflavine nanocapsules on choroidal neovascularization

Following confirmation of free acriflavine’s ability to inhibit CNVM formation *in vivo*, we proceeded with testing of acriflavine nanocapsules. Each animal received five laser spots in both eyes, followed by a single intravitreal injection of either blank nanocapsules, acriflavine nanocapsules, free acriflavine, or PBS in the right eye. The contralateral eye was left untreated and used as a disease only control. From our previously published drug release studies (*29*) we noted that approximately 25% of the drug is released from the nanocapsules in the span of two weeks, and as such we used ∼600 ug of acriflavine capsules, containing 30ug of acriflavine. At 14-days following laser injury, CNVM area was evaluated (**Figure 4**). As expected, nanocapsules were clearly visible via both fluorescent angiography (**Figure 4B**) and fundus photography (**Figure 4J**). No evidence of clinical complications, including ocular inflammation or cataract formation, was detected in any animal. To evaluate CNVM size RPE/choroid flat mounts were labeled with GSL-1 (**Figure 4E-H**). Animals in both free acriflavine and acriflavine nanocapsule groups had significantly smaller CNVMs compared to drug free controls (**Figure 4I**). While there was no significant difference in CNVM area between the free acriflavine and acriflavine nanocapsule conditions (**Figure 4I**), the number of CNVMs present at 14-days following laser injury was reduced in animals that received acriflavine nanocapsules. Specifically, approximately 60% of the laser spots formed CNVMs in the acriflavine nanocapsule group, compared to 76% in the free acriflavine group, and 84% in the drug free control group.

**Figure 4.**
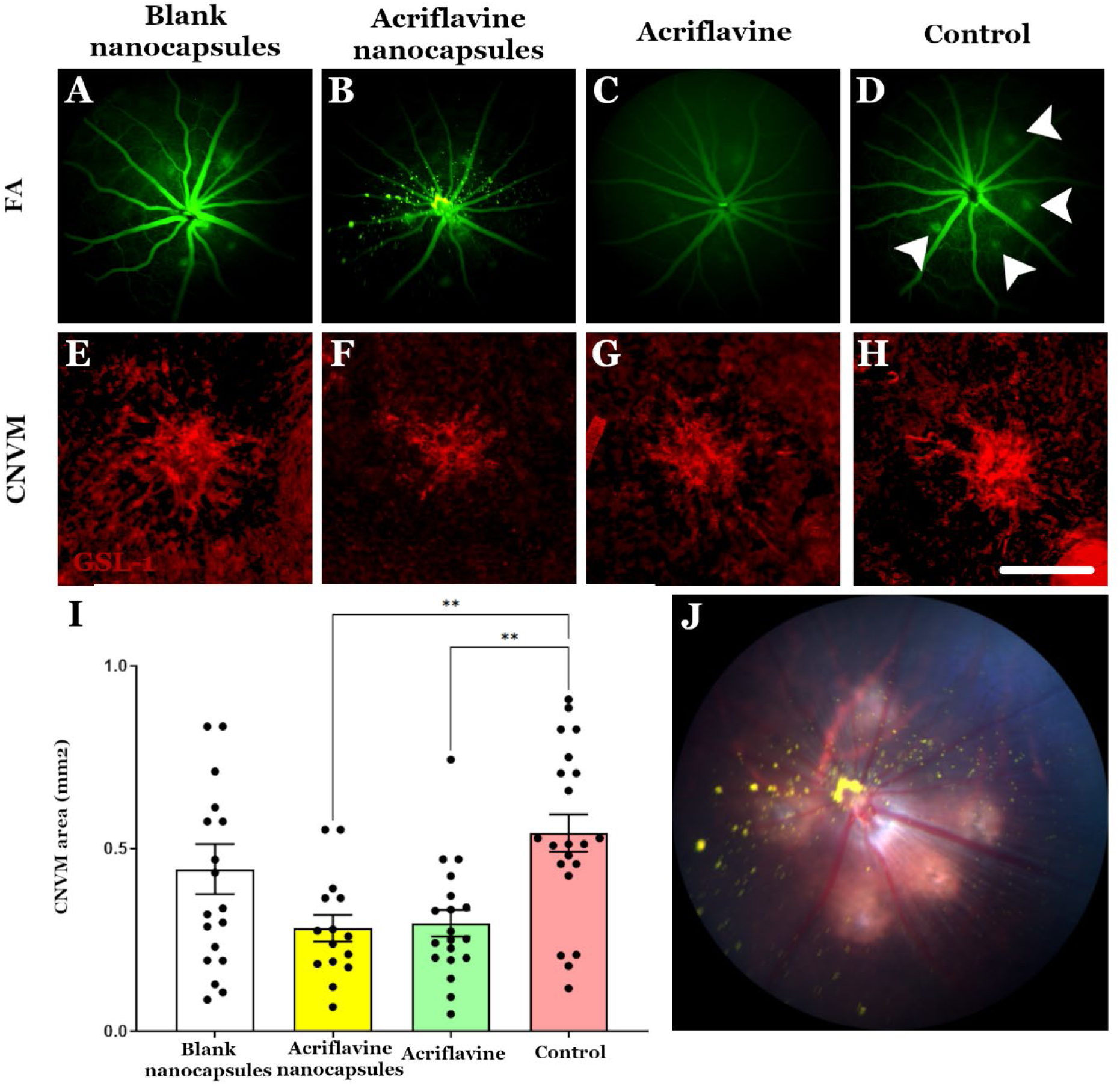
Nanocapsule delivered acriflavine inhibits CNVM formation. A-D) Fluorescence fundus angiographs (FA) showing CNVMs (e.g., arrowheads in panel D) in each of the different treatment groups at 14-days post-laser injury. On day-0, all the animals received 5 laser spots along with an intravitreal injection of either PBS, free acriflavine, blank nanocapsules, or acriflavine loaded nanocapsules. **E-H)** Immunofluorescence imaging of the flat mounted choroid-sclera complex labeled with GSL1 (*Griffonia simplicifolia* lectin I) to assess CNVM area (Scale bar is 1000 µm). **I)** CNVM area in each of the different treatment groups at 14-days following laser photocoagulation (** indicates a p-value <0.01, obtained from one way ANOVA, using Tukey’s multiple comparison test). **J)** Fundus photo of a Brown Norway rat that received a dose of acriflavine loaded polyurethane nanocapsules (yellow, suspended in the vitreous).

As acriflavine is a small molecule that can pass through the blood retinal barrier, we next wanted to determine if treatment had an impact on CNVM formation in the contralateral eye (i.e., the left eye of each animal received 5 laser spots without intravitreal drug delivery). There was no significant difference in the area or number of CNVMs formed in the contralateral eye of animals in the control vs. treatment groups (**Supplementary Figure 1A**).

To examine CNVM morphology and determine impact on retinal architecture using histology, the experiment was repeated, eyes were dissected, sectioned, and stained using the tomato lectin from *Lycopersicon esculentum* (LEL) (**Figure 5**). CNVM morphology and damage to the overlying neural retina were strikingly different between treatment versus control eyes. For instance, in PBS (**Figure 5A**) and blank nanocapsule (**Figure 5C**) control groups, the neural retina, including the inner nuclear layer, above the CNVM was severely disrupted. In some cases, CNVMs extended not only intraretinally (**Figure 5C**) but through the retina into the vitreous (**Figure 5A**). While the neural retina above the CNVMs in the treatment groups was not normal, damage was largely limited to the outer nuclear layer, which showed undulating folds with an intact layer of photoreceptors. For instance, in both the free acriflavine (**Figure 5B**) and acriflavine nanocapsule groups (**Figure 5D**) the CNVM was restricted to the subretinal space between the RPE and photoreceptor cell layers.

**Figure 5.**
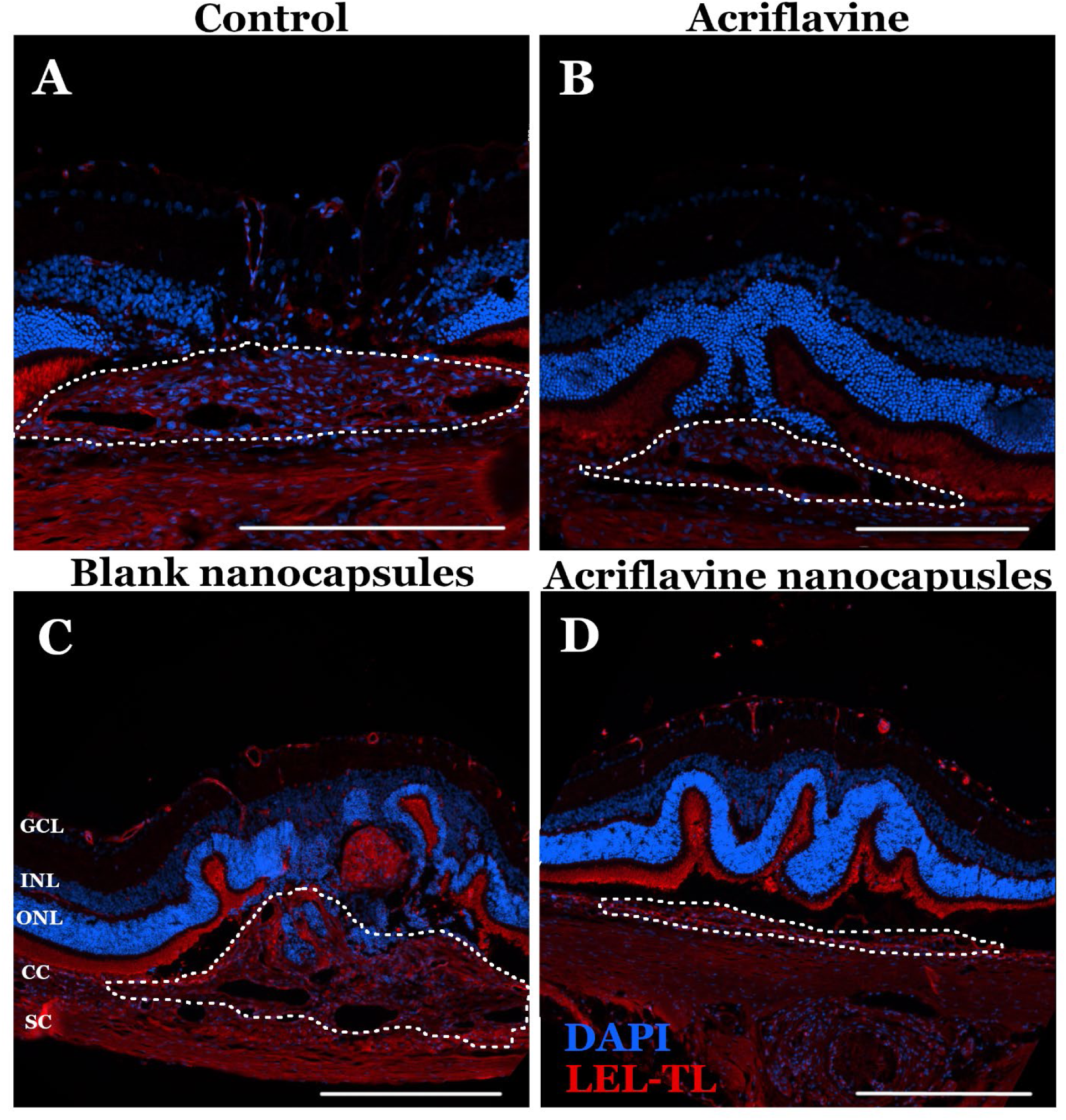
Acriflavine nanocapsules reduce CNVM size and protect the overlying neural retina. A-D) Immunofluorescence staining of rat retina using the tomato lectin LEL at 14-days following laser injury and intravitreal injection of PBS (**A**), free acriflavine (**B**), blank nanocapsules (**C**), and acriflavine nanocapsules (**D**). Scale bar = 200μm.

### Comparison of the impact of acriflavine by different routes of administration

In a recent study by Hackett and colleagues, suprachoroidal delivery of acriflavine loaded PLGA microparticles was found to be superior to intravitreal delivery for CNVM treatment(*26*). To determine if route of delivery had a similar impact on treatment efficacy when delivered by nanocapsules, CNVMs were induced, and acriflavine loaded polyurethane nanocapsules (in 60 mg/ml; 10ul per eye) were delivered via intravitreal, subretinal, or suprachoroidal injection. While there was a small but non-significant reduction in CNVM area in eyes that received acriflavine nanocapsules via both suprachoroidal and subretinal injection (**Figure 6A**), fundus photography revealed widespread retinal atrophy in these groups (**Supplementary Figure 2**). In the intravitreal injection group retinal injury was confined to the injection site (**Figure 6B**). Retina outside of the injection site was intact and lacked immune cell infiltration and Muller glial cell activation (**Figure 6C**). Conversely, suprachoroidal (**Figure 6D**) and subretinal injection (**Figure 6E**) caused significant retinal damage marked by ONL thinning, rosette formation and glial cell activation.

**Figure 6.**
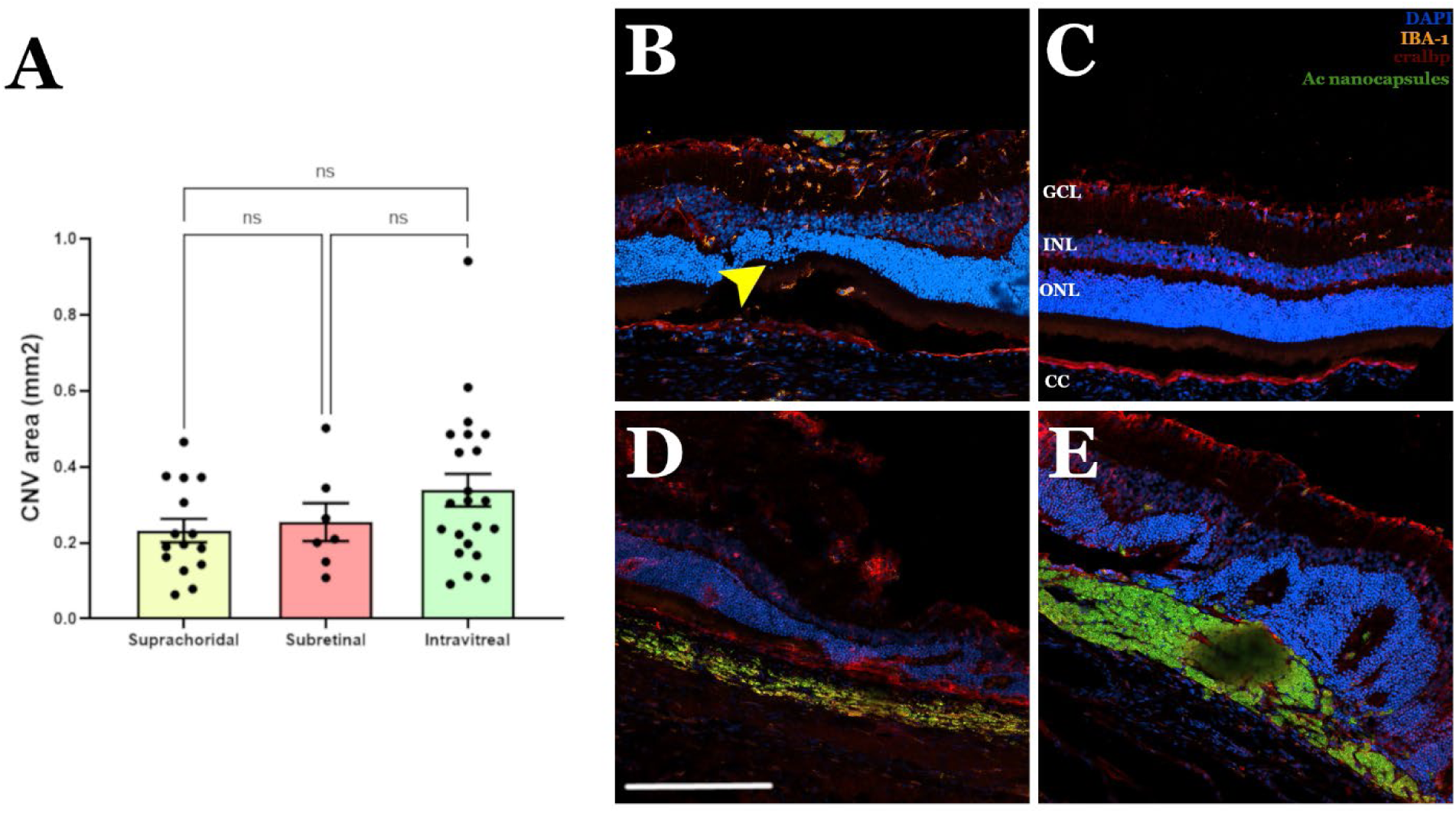
Impact of ocular delivery route on treatment efficacy. **A)** Histogram depicting CNVM area at 14-days following laser injury and suprachoroidal, subretinal, or intravitreal injection of acriflavine nanocapsules (data obtained from GSL1 labeled RPE/choroidal flat mounts, each data point represents the area of one CNVM formed; each animal received 5 laser spots per eye followed by injection (n=6)). Immunohistochemical analysis at 14-days following laser injury and intravitreal **(B,C)**, suprachoroidal **(D)** or subretinal **(E)** injection of acriflavine nanocapsules. While suprachoroidal and subretinal injection of acriflavine nanocapsules induced a non-significant reduction in total CNVM area, both routes of delivery resulted in widespread glial cell activation, immune cell infiltration, and retinal injury. Anti-IBA1 was used to label infiltrating immune cells and resident microglia. DAPI was used as a nuclear counterstain. Arrowhead indicates auto fluorescent polyurethane nanocapsules. Scale bar = 200 μm

## Discussion

Despite the overwhelming therapeutic success of anti-VEGF agents for the treatment of ocular neovascularization, alternate therapeutic targets with better effect or longer durability of drug effect remain needed. Endogenously expressed VEGF is required for maintenance of the neural retina (*16, 17*) and in some cases over time neovascular membranes become intractable to anti-VEGF treatments (*35, 36*). Acriflavine, an FDA-approved small molecule inhibitor of HIF-1α (a master regulator of angiogenesis), is known for its antiseptic, antibacterial, and trypanocidal properties (*24*). In addition to inhibiting the HIF-1α pathway, acriflavine has also been shown to inhibit activation of nuclear factor-kappa B (NF-κB) and production of tumor necrosis factor-α (TNF-α). Blocking NF-κB and TNF-α contribute to acriflavines reported anti-inflammatory potential (*37, 38*), an added benefit given that repeated intravitreal injection may be associated with an increased risk of ocular inflammation (*14, 39*).

Polyurethanes are highly tunable polymeric compounds with organic units joined by urethane (carbamate) linkages (*40*). They have been used for a variety of different applications ranging from heart valves to pacemakers, vascular grafts, catheters, and drug delivery vehicles (*41–44*). Polyurethanes can be fabricated as transparent materials with UV absorbing properties that have excellent biocompatibility, which makes them promising candidates for ophthalmological applications. In fact, they have been used for several ocular indications including intraocular lenses, keratoprostheses, and wound closures for corneal perforations (*45–48*). More recently, we have used polyurethane to fabricate nanocapsules for sustained ocular drug delivery (*29, 41*). While it can be challenging to encapsulate hydrophilic drugs such as acriflavine within hydrophobic polyurethanes, using a water and oil emulsion strategy, we successfully achieved a 5% loading rate, which is well within the typical range for nanocapsule drug delivery systems with long-term release kinetics (*29, 49*). To decrease their hydrophobicity and enabled suspension in aqueous buffers for intraocular injection (e.g. sterile saline), we coated nanocapsules with PEG and poloxamer. Following production, Acriflavine loaded polyurethane nanocapsules were found to have a biphasic release profile. An initial burst characterized by release of approximately 20% of encapsulated drug, and sustained deaccelerated delivery, with 30% of encapsulated drug released over the following 90 days (*29*). In this study, we demonstrate that this release profile was superior to injection of free drug as a single bolus injection for CNVM treatment. Specifically, in addition to reducing CNVM area, sustained release prevented CNVMs from disrupting continuity of the retinal outer nuclear layer and destroying photoreceptor cell inner and outer segments, a phenomenon that was seen in both negative controls and the free acriflavine condition (**Figure 5**).

While not exploited in this study, an added benefit of polyurethane nanocapsules is the ability to trigger on demand drug release using noninvasive ultrasound (*29*). Specifically, we previously demonstrated that 0.5ug/mg of acriflavine (1% of encapsulated drug) was released per each 20 second ultrasound pulse given (*29*). If despite sustained release, which we showed here was highly effective, a patient did develop a CNVM the ability to noninvasively stimulate bursts of drug release via ultrasound could enable treatment in a local eye clinic, negating the need for rural patients to drive long distances to tertiary care facilities.

While intravitreal injection is the most common route of anti-VEGF agent delivery, as indicated above this has been associated with complications such as intraocular inflammation and infectious endophthalmitis (*14, 39*). To address this concern, Hackett and colleagues recently evaluated suprachoroidal delivery of acriflavine loaded PLGA microparticles (*26*). As compared to intravitreal injections, they found that suprachoroidal injection further reduced neovascularization. While not statistically significant, we found a similar trend in this study. Specifically, there was a slight decrease in CNVM area in animals that received acriflavine nanocapsules via both subretinal and suprachoroidal injection as compared to those that were given intravitreal injections. Unfortunately, this slight improvement in treatment effect was overshadowed by the negative impact that suprachoroidal and subretinal injection had on retinal anatomy. While the neural retina remained intact in the intravitreal group, animals that received the subretinal and suprachoroidal injections had significant damage to both the RPE and neural retina **(Figure 6, Supplementary Figure 2**). As placement of polyurethane beads in the subretinal space physical separates the RPE and photoreceptor cell layers, the fact that this mode of delivery was poorly tolerated was somewhat expected. That said, given the previous report by Hackett and colleagues that suprachoroidal injection of PLGA based microbeads was superior to intravitreal injection, we were surprised to find that suprachoroidal delivery of polyurethane nanocapsules in this study was so disruptive. While the reason for this difference is unknown, it may in part be due to the difference in release kinetics afforded by polyurethane versus PLGA. Specifically, it is possible that the amount of acriflavine being consistently released from our nanocapsule formulation is higher than that of the PLGA particles fabricated in by Hacket et. al., and that prolonged sustained delivery of high doses of acriflavine to the uninjured choroid disrupts the normal vasculature and overlying retina. Alternatively, it could simply be a result of injection trauma, namely a byproduct of the mechanical forces caused by hydraulic dissection of the suprachoroidal space in the rat.

In conclusion, in this study we successfully demonstrate fabrication and intraocular deliver of acriflavine loaded nanocapsules in a rat model of choroidal neovascularization. Nanocapsule mediated sustained acriflavine delivery reduced neovascular membrane size, decreased neovascular membrane occurrence, and prevented destruction of the overlying neural retina. While CNVM area was reduced further following subretinal and suprachoroidal injection, these methods of delivery were poorly tolerated. Future studies may evaluate the impact of clinically relevant surgical delivery methods and human equivalent drug doses on retinal anatomy and function through testing in large animal preclinical models such as the pig.

## Methods

### Polyurethane nanocapsule synthesis

Polyurethane nanocapsules were synthesized using our previously described protocol (*29*). In brief, 1.15 ml of hexadecane (Fischer Scientific, Waltham, MA) was added to 70 ml of solution of 1% sodium dodecyl sulfate (Fischer Scientific, Waltham, MA) in water and stirred for 1 hour at 40°C. Then a solution of 2.1 ml of isophorone diisocyanate (Fischer Scientific, Waltham, MA) and 110 mg of acriflavine (Sigma Aldrich, St. Louis, MO) in 7 ml of water was added dropwise to the first solution. The mixture was emulsified by sonicating at 38% amplitude for 1 min using a 500W probe sonicator (QSonica, Newtown, CT). A solution of 5.9 gm of hexanediol (Fischer Scientific, Waltham, MA) in 10ml water was added to the emulsion while sonicating and followed by a solution mPEG-OH (1 gm in 10 ml; 5kDa; CreativePEGworks, Chapel Hill, NC). This emulsion was stirred at 40°C and 300 rpm for 24 hours. The resulting nanocapsules were separated by ultracentrifugation at 10,000 rpm for 10 mins. The nanocapsules were subsequently washed by resuspending in a 1% Poloxamer 188 PRO (mol wt. 8400 Da, Sigma Aldrich, St. Louis, MO) solution in water for 5 mins, followed by ultracentrifugation. This was repeated three times and after the final spin the nanocapsules were resuspended in 5 ml of ultrapure water, flash frozen and lyophilized.

### Nanocapsule loading and sizing

To determine drug loading, the polyurethane nanocapsules were dissolved in ethanol and their fluorescence was measured across a standard curve of acriflavine in ethanol (excitation - 426 nm; emission - 524 nm) (*50*). Nanocapsules were dispersed in PBS at ∼25 µg/ml and characterized for size using a Multi-laser nanoparticle tracking analyzer (ViewSizer3000, Horiba, Kyota, Japan). SEM images were obtained using Hitachi S-4800 (Hitachi, Tokyo, Japan)

### Human choroidal endothelial cell culture and scratch assay

Choroidal endothelial cells (ICEC2-TS) were cultured in complete endothelial cell growth media (R&D systems) supplemented with 1% primocin (InvivoGen, San Diego, CA). For the scratch assay, 200,000 cells were seeded per well in a 6 well tissue culture plate and grown until confluent. A scratch was made along the diameter of the well using a 200 μl universal pipette tip and the requisite amount of acriflavine or nanocapsules were resuspended in complete media and added to the respective wells. Wells are imaged at different time points using BioTek Cytations 5 (Agilent Technologies, Santa Clara, CA).

### Laser injury and ocular injection

6-8 week old Brown Norway rats (Charles River, Wilmington, MA), equal number of males and females for each cohort (n=6), were used for all *in vivo* studies. Animals were anesthetized by intraperitoneal injection of ketamine (91 mg/kg) and xylazine (9.1 mg/kg) and their eyes were dilated using 1% tropicamide (Alcon Laboratories, Fort Worth, TX). Laser photocoagulation was performed using a PASCAL argon green laser (OptiMedica, Sunnyvale, CA) at 150mW power, and 100-micron spot size with an exposure time of 10 msec.

Ocular injections were performed as previously described (*51*) immediately following laser photocoagulation. Briefly, under direct visualization using an operating microscope, a limited temporal conjunctival peritomy was created. A 30-gauge needle was used to create a slit in the sclera, and a 33-gauge blunt-tipped Hamilton syringe (Hamilton Company, Reno, NV) was used to inject product into the vitreous cavity, subretinal space, or suprachoroidal space. Eyes were examined immediately post-injection to confirm successful injections and absence of surgical complications. Nanocapsules (60 mg/ml) and free drug (0.75 mg/ml) were suspended in sterile PBS and sonicated for 2 mins in a sonicating water bath prior to use. All injections were 10 μl total volume.

### Fundus photos and Fluorescein angiography

Animals were examined 14-days post-injection. Prior to sacrifice, eyes were examined under an operating microscope to record clinical findings, including any complications such as inflammation, cataract formation, or retinal detachment. Optical coherence tomography (OCT) was performed (OCT; Phoenix Micron Image-Guided OCT2, Phoenix Laboratories, Pleasanton, CA) to evaluate the CNV formation. A 10% solution of Fluorescein (Sigma Aldrich, St. Louis, MO) was injected intravitreally to obtain angiograms using a rodent-specific fluorescent fundus camera (Micron IV, Phoenix Laboratories, Pleasanton, CA).

### Tissue processing, imaging and analysis

Following clinical imaging, animals were euthanized using euthasol (150 mg/kg). Eyes were subsequently enucleated and fixed in 4% PFA in PBS at 4C for 5 hours. Cornea, lens and the iris were dissected to obtain the posterior segment. For analysis of retinal anatomy eyes were cryopreserved, frozen, sectioned and labeled with anti-CRALBP(1:400; Abcam, San Diego, CA), anti-IBA1 (1:400; Fujifilm Wako Chemicals, Richmond, VA), and Tomato lectin(1:50; Vector Laboratories, Burlingame, CA). DAPI was used as a nuclear counterstain. For flat mounting, the retina was peeled off from the posterior segment. The remaining choroid-scleral complex was incubated with Dylight594 conjugated GSL-1 (1:50 dilution; Vector Laboratories, Burlingame, CA) overnight at 4°C on a rocker. Then the tissue is washed, cloverleafed via cutting at 90-degree intervals and mounted onto a glass slide with cover slip. The CNVs formed were imaged using a fluorescence microscope (BZ-X810, Keyence, Osaka, Japan) and analyzed using Fiji (*52*).

## Acknowledgements

We would like to thank the University of Iowa Animal Care staff for their help and commitment to ensuring the success of animal experiments.

