## Supplementary for "Acriflavine delivery via Polyurethane nanocapsules to treat neovascular age-related macular degeneration"

**Funding:** NIH R01 EY035676

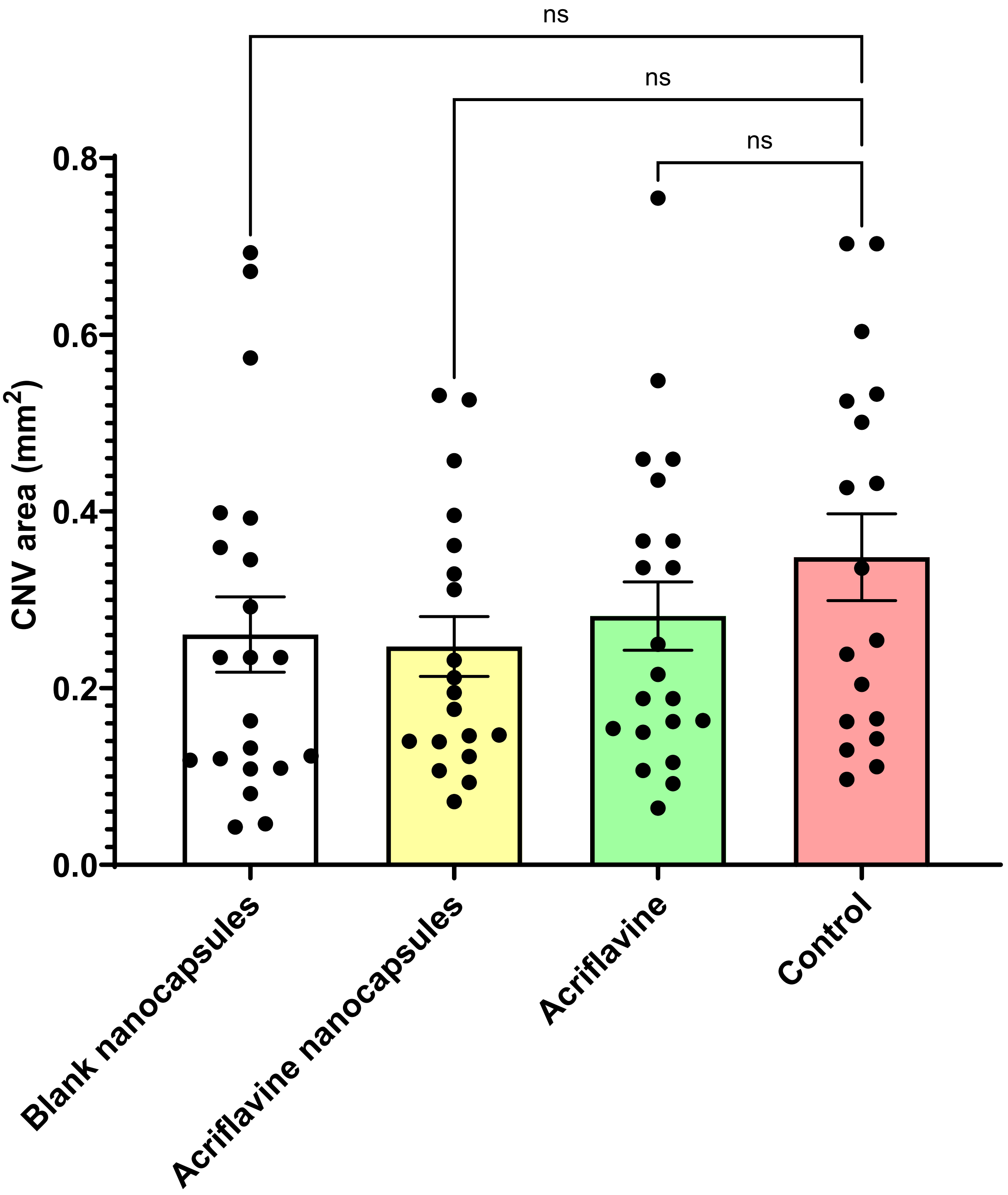


**Supplementary Figure 1.** Size of CNVs formed in the left eye (un injected) of the animals that received different treatments via intravitreal injections in the right eye

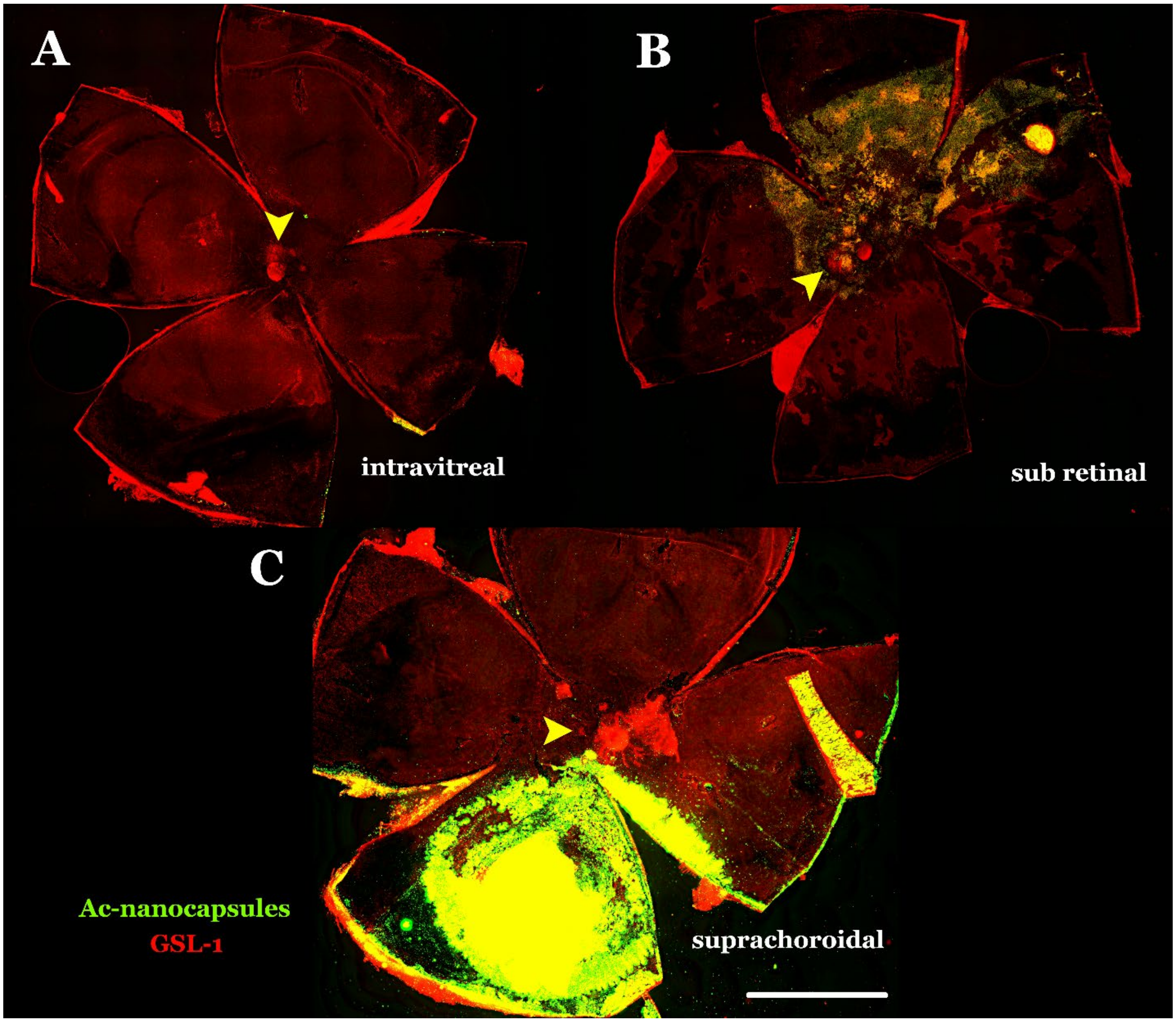


**Supplementary Figure 2.** RPE choroid flat mounts showing formation of CNVs (around the optic nerve indicated by yellow arrows) stained by GSL-1 red and acriflavine nanocapsules (green) in different animals that received these nanocapsules via different routes A) intravitreal B) sub retinal and C) suprachoroidal (Scale bar is 1000 µm)
